# Genetic dissection of assortative mating behavior

**DOI:** 10.1101/282301

**Authors:** Richard M. Merrill, Pasi Rastas, Simon H. Martin, Maria C. Melo, Sarah Barker, John Davey, W. Owen McMillan, Chris D. Jiggins

**Affiliations:** Division of Evolutionary Biology, Ludwig-Maximilians-Universität, München, Germany; Department of Zoology, University of Cambridge, Cambridge, UK; Smithsonian Tropical Research Institute, Panama City, Panama; IST Austria, Klosterburg, Austria; Department of Biology, University of York, York, UK

## Abstract

The evolution of new species is made easier when traits under divergent ecological selection are also mating cues. Such ecological mating cues are now considered more common than previously thought, but we still know little about the genetic changes underlying their evolution, or more generally about the genetic basis for assortative mating behaviors. Both tight physical linkage and the existence of large effect preference loci will strengthen genetic associations between behavioral and ecological barriers, promoting the evolution of assortative mating. The warning patterns of *Heliconius melpomene* and *H. cydno* are under disruptive selection due to increased predation of non-mimetic hybrids, and are used during mate recognition. We carried out a genome-wide quantitative trait locus (QTL) analysis of preference behaviors between these species and showed that divergent male preference has a simple genetic basis. We identify three QTLs that together explain a large proportion (∼60%) of the differences in preference behavior observed between the parental species. One of these QTLs is just 1.2 (0-4.8) cM from the major color pattern gene *optix*, and, individually, all three have a large effect on the preference phenotype. Genomic divergence between *H. cydno* and *H. melpomene* is high but broadly heterogenous, and admixture is reduced at the preference-*optix* color pattern locus, but not the other preference QTL. The simple genetic architecture we reveal will facilitate the evolution and maintenance of new species despite on-going gene flow by coupling behavioral and ecological aspects of reproductive isolation.

## Introduction

During ecological speciation, reproductive isolation evolves as a result of divergent natural selection [1]. Although ecological barriers can reduce gene flow between divergent populations, speciation normally requires the evolution of assortative mating [1,2]. This is made easier if traits under divergent ecological selection also contribute to assortative mating, as this couples ecological and behavioral barriers [3–6]. Ecologically relevant mating cues (sometimes known as ‘magic traits’ [2,6]) are now predicted to be widespread in nature [6,7], and the last few years have seen considerable progress in our understanding of their genetic basis. For example, studies have explored the genetic basis of beak shape in Darwin’s finches [8], body shape in sticklebacks [9,10], cuticular hydrocarbons in *Drosophila* [11], and wing patterns in *Heliconius* butterflies [12–14]. However, the extent to which these traits contribute to assortative mating depends on the evolution of corresponding preference behaviors, and the underlying genetic architecture.

We still know little about the process by which ecological traits are co-opted as mating cues, and in particular, how matching cues *and* preference behaviors are controlled genetically (but see [15]). Both the substitution of large effect preference alleles, and physical linkage will strengthen linkage disequilibrium (‘LD’, *i.e.* the non-random association of alleles at different loci [16]) between cue and preference. Strong LD between barrier loci is expected to both maintain and facilitate the evolution of new species in the face of gene flow. This is the result of two key, but related processes. First, LD between barrier loci will result in the coupling of barrier effects, and where these effects coincide the overall barrier to gene flow is increased [4,16]. Second, LD between pre-and post-mating barrier loci will facilitate an increase in premating isolation in response to selection against hybridization (*i.e.* reinforcement, *sensu* [18]), by transferring the effects of selection from the latter to the former [19].

In central Panama, the butterfly *Heliconius melpomene rosina* is a precise Müllerian mimic of *H. erato* and normally occurs in forest-edge habitats, whereas the closely related species *H. cydno chioneus* mimics *H. sapho* and is more common in closed-forest habitats, although *H. melpomene* and *H. cydno* are often seen flying together (Fig. 1a& b) [20]. Hybrids are viable but occur at very low frequency in the wild (estimated at ∼0.1%), consistent with strong assortative mating shown in insectary experiments. Specifically, heterospecific mating was not observed in 50 choice and no-choice trials between Panamanian *H. melpomene* and *H. cydno* ([21,22] ; see also [23]).

**Figure 1.**
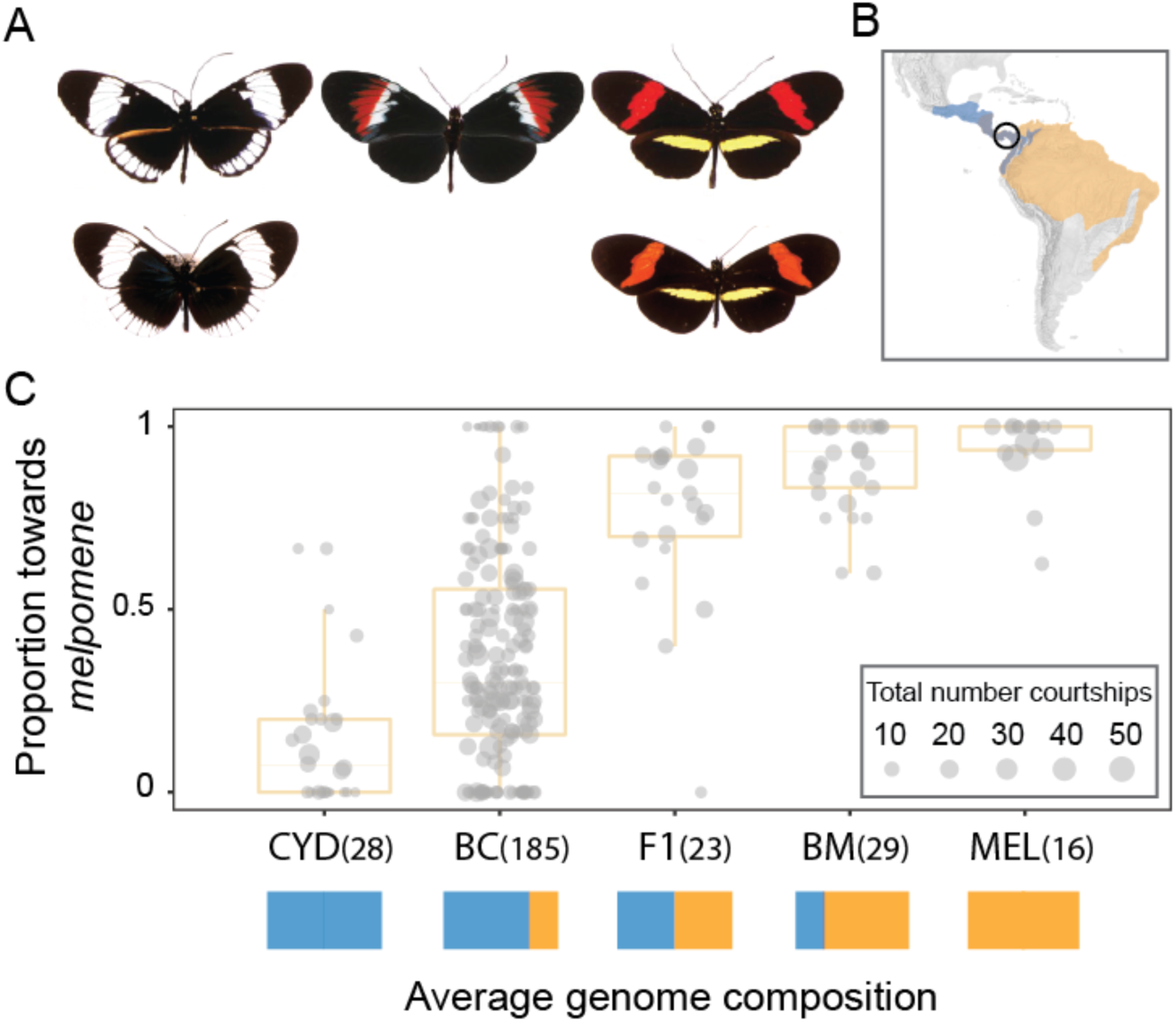
Divergence in warning pattern cue and corresponding preference in sympatric *Heliconius* butterflies. A, Wing pattern phenotypes of: top, *Heliconius cydno chioneus* (left), *H. melpomene rosina* (right) their non-mimetic first generation hybrid (center), and bottom, their sympatric co-mimics *H. sapho sapho* (left) and *H. erato demophoon* (right). B, Distribution of *H. cydno* (blue) and *H. melpomene* (orange). Individuals were collected, and experiments performed in Panama (black circle), where the two species co-occur in Central and northern South America. **C**, Proportion of courtships directed towards *H. melpomene* (as opposed to *H. cydno*) females for *H. cydno* (CYD), *H. melpomene* (MEL), their F1 and backcross hybrids to *H. cydno* (BC) and *H. melpomene* (BM). Values in parentheses indicate total number of individuals with behavioral data. Solid colored boxes represent expected average genome contribution of each generation. Note that a further 11 BC individuals were tested but performed no courtship behaviors.

The amenability of *Heliconius* color patterns to experimental manipulation has led to the demonstration that color pattern is both under strong disruptive selection due to predation [24], and also that males prefer live females and paper models with the same color pattern as themselves [24]. These results led Servedio and colleagues [6] to conclude that, unlike other putative examples, both criteria for a magic trait have been confirmed with manipulative experiments in *H. melpomene rosina* and *H. cydno chioneus*. Although female preferences undoubtedly contribute to assortative mating [25–27], male preferences act first in these species such that strong observed male discrimination against heterospecific females will have a disproportionate contribution to overall reproductive isolation [28]. As highlighted by Coyne and Orr [29], the order in which reproductive isolation acts influences their relative contribution to overall isolation. In this case, the ordering of behavioral decisions is likely predetermined by their sensory systems: *Heliconius* lack specialized olfactory structures to support long range detection of chemical signals, so are only likely to use these in close proximity, whereas they have very good long-range vision [30]. As such, not only is male preference in *Heliconius* butterflies experimentally more tractable than other components of behavioral isolation, it is also an important barrier to gene flow.

Crossing experiments have shown that the shift in mimetic warning pattern between *H. melpomene rosina* and *H. cydno chioneus* is largely controlled by just three major effect loci [31]. Genes underlying these loci have now been identified: the transcription factor *optix* controls red patterns [12], the *WntA* gene controls forewing band shape [13] and yellow patterns map to the gene *cortex* [14]. In addition, a further locus, *K*, segregates in crosses between *H. melpomene rosina* and *H. cydno chioneus* with more minor effect [31]. Further modularity occurs within these loci. For example, different regulatory elements of *optix* each result in distinct red pattern elements [32]. The modular nature of individual color pattern loci and their functionally sufficient enhancers means that they can be combined to produce considerable phenotypic diversity [32,33]. These loci are large-effect ‘speciation genes’, in that they control traits that generate strong reproductive isolation [34].

Two of these color pattern loci, *optix* and *K,* have previously been associated with *Heliconius* courtship behaviors [25,35,36]; however, these studies do not provide evidence for tight physical linkage (<20cM) between warning pattern and preference loci. Our own previous study tested for an association between Mendelian color pattern loci and preference behaviors [25], but did not correct for the segregation of alleles across the genome, so that reported levels of support are likely inflated [37]; and an earlier study of the parapatric *taxa H.* cydno *and H. pachinus* [35] is limited by small sample size [37]. Regardless of the level of statistical support for preference QTL, these studies both lack the resolution to demonstrate the degree of tight physical linkage between loci contributing to reproductive isolation that would be expected to aid speciation. Perhaps the best evidence comes a study of wild *H. cydno alithea* [36]. This population is polymorphic for a yellow or white forewing (due to the segregation of alleles at the *K* locus), and males with a yellow forewing prefer yellow females. These results are important because they suggest a key component of speciation: Specifically, coupling between potential behavioral and ecological barriers. However, because they rely on segregation within a wild population, rather than laboratory crosses, it is not possible to distinguish physical linkage from genetic associations between cue and preference alleles due to non-random mating. The extent to which warning pattern and behavioral loci are physically linked in *Heliconius*, as well as the existence of major preference loci elsewhere in the genome remains unknown. To address this, and to complement our extensive knowledge of the genetics of their color pattern cues, here we use a genome-wide quantitative trait locus (QTL) approach to explore the genetics of male preference behaviors between the sympatric species *H. melpomene rosina* and *H. cydno chioneus*.

## Results

We studied male mating preference among F1 and backcross hybrid families between *H. melpomene rosina* and *H. cydno chioneus*, in standardized choice trials [25,38] (Figs. 1 and S1). We introduced individual males into an experimental cage and recorded courtship directed towards two virgin females, one of each species. In total, we collected data from 1347 behavioral trials, across 292 individuals. Multiple trials were performed for each backcross male, from which we determined the relative courtship time directed towards *H. melpomene* and *H. cydno* females.

### Three loci contribute to species differences in preference behavior

As reported previously [25], F1 males have a strong preference for the red *H. melpomene* females, and little segregation in mate preference is observed among the backcross to *melpomene* (and whose mean preference does not differ significantly from that of pure *H. melpomene* males: 2ΔlnL = 1.33, *d.f.* = 1, *P* > 0.2), implying that *melpomene* mate preference alleles are dominant. In contrast, courtship behavior segregates among *H. cydno* backcross males, permitting analysis of the genetic basis for this mating behavior (Fig. 1C). Consequently, all subsequent analyses were performed on backcross to *cydno* males. We used a genome-wide quantitative trait locus (QTL) mapping approach to identify the genomic regions underlying divergence in mate attraction. Linkage maps were constructed from genotype data of 331 backcross-to-cydno individuals and their associated parents [39], including 146 individual males for which we had recorded attraction behaviors.

We identified three unlinked QTLs on chromosomes 1, 17 and 18 associated with variation in the relative amount of time males spent courting red *H. melpomene* and white *H. cydno* females (Fig. 2A). Of these, one is tightly linked to the *optix* locus on chromosome 18, which controls the presence/absence of a red forewing band. Specifically, the QTL peak for the behavioral QTL on chromosome 18 (at 0cM) is just 1.2cM from *optix.* The associated 1.5-LOD support interval is between 0 and 6.0cM, suggesting that the true location of the QTL is no more than 4.8cM from the *optix* coding region (whose genetic position is at 1.2cM) (Fig. 3); however, given that the peak support (i*.e.* highest LOD score) for our behavioral QTL is at 0cm and that this rapidly drops off, physical linkage between wing patterning cue and preference loci is likely much tighter than a strict 1.5-LOD interval might suggest. In contrast, the QTL on chromosome 1 is at least 30cM from the gene *wingless*, which although unlikely to be a color pattern gene itself has previously been associated with the *K* wing pattern locus between taxa within the *cydno* clade [35]. No known wing pattern loci reside on chromosome 17 and this chromosome does not explain any of the pattern variation segregating in our BC pedigrees (Merrill, unpublished data).

**Figure 2.**
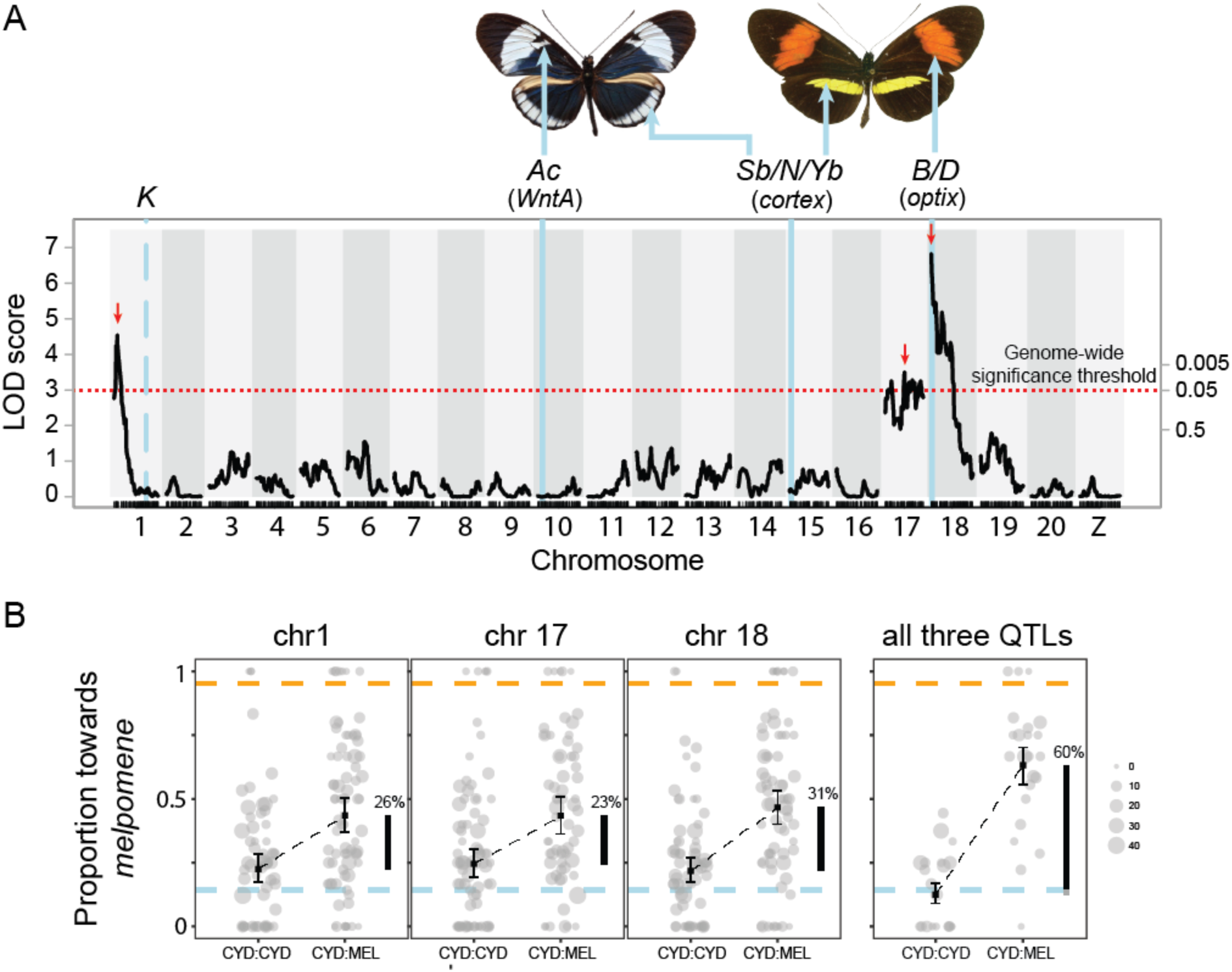
QTL analysis of variation in mate preference. A, QTLs for relative time males court *H. melpomene* (as opposed to *H. cydno*) females on chromosomes 1, 17 and 18 (*n* = 139). Scale on right axis depicts genome-wide significance, determined through permutation, corresponding to the LOD score as shown on the left axis. Dotted red line represents log odds ratio (LOD) significance threshold (genome-wide alpha = 0.05, LOD = 2.99). Dashes indicate position of genetic markers (SNPs) and red arrows indicate the position of the max LOD score for each QTL (used in B). Vertical blue lines represent the position of major color pattern loci, and their phenotypic effects. Note that the *K* locus only has limited phenotypic effects in crosses between *H. cydno chioneus* and *H. melpomene rosina*, but is responsible for the switch from yellow to white color pattern elements between other taxa within the *melpomene-cydno* clade. B, Proportion of time males court *H. melpomene* (as opposed to *H. cydno*) females for each of the two genotypes for respective QTLs (homozygous = CYD: CYD, and heterozygous = CYD:MEL). Error bars represent 95% confidence intervals. Lower dashed blue and upper orange bars represent mean phenotypes measured in *H. cydno* and *H. melpomene,* respectively. Circle size depicts total number of ‘courtship minutes’ for each male. Vertical black bars indicate the percentage of the difference measured in the parental species explained.

**Figure 3.**
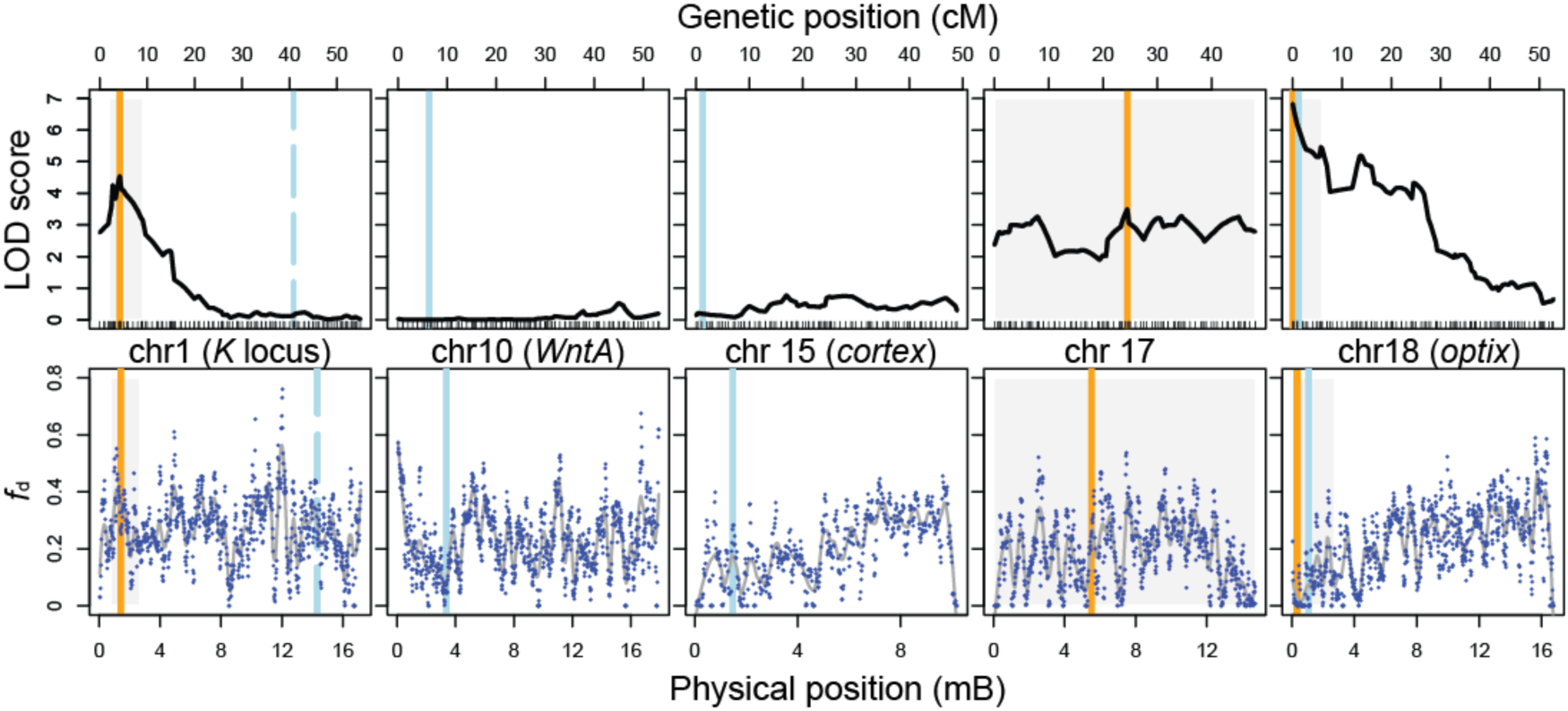
Genetic and physical positions of behavioral QTL and the warning pattern loci, and localized levels of admixture (*f*_d_). Vertical blue lines represent the position of major color pattern loci and orange lines represent the position of peak LOD score for each behavioral QTL. Gray boxes indicate the 1.5-LOD support interval for each QTL. Top panel: Dashes along the x-axis indicate position of genetic markers (SNPs). Bottom panel: Blue points represent *f*_d_ values for 100kb windows. *f*_d_ was measured between *H. melpomene rosina* and *H. cydno chioneus* individuals from populations samples in Panama; *H. melpomene melpomene* from French Guiana, which is allopatric with respect to *H. cydno*, was the ‘control’ population.

Modeling supports additive effects of all three detected loci (Table 1), and in our mapping population these three QTLs together explain ∼60% of the difference in male preference behavior between the parental species (Fig. 2B). Given the sample sizes feasible in *Heliconius*, our analysis lacks the power to resolve smaller effect QTLs. We also found no evidence of pairwise interactions between individual QTLs in our model of relative courtship time, which again is unsurprising given achievable sample sizes. However, genome scans considering individuals with alternative genotypes at the QTL on chromosome 18 separately revealed a significant QTL on chromosome 17 (LOD = 3.52, *P*=0.016) for heterozygous (i.e. LG18@0cM = CYD:MEL), but not for homozygous (i.e. LG18@0cM = CYD:CYD) males (Fig. S2), though this result is not supported by non-parametric interval mapping (LOD = 2.4, *P*=0.132). Nevertheless, these results perhaps suggest that alleles on chromosomes 17 and 18, or the specific behaviors they influence, may interact.

**Table 1.** Individual and combined QTLs for differences in relative courtship time.

| Chromosome | Position (cM) | LOD score | $\Delta p\text{LOD}_a$ | $2\Delta\ln L$ | $P$ |
| --- | --- | --- | --- | --- | --- |
| 1 | 4.2 (2-9.1) | 4.54 | -1.04 | 18.54 | <0.001 |
| 17 | 24.4 (0-48.3) | 3.50 | -1.03 | 18.50 | <0.001 |
| 18 | 0 (0-6) | 6.83 | -3.87 | 31.60 | <0.001 |
| **1+17+18** | **–** | **14.90** | **(5.93)** | **–** | **–** |

Position in cM (1.5 LOD interval); LOD score, log odds ratio; $\Delta p\text{LOD}_a$ , penalized LOD score, change in penalized LOD score compared to the full (best supported) model incorporating all three putative QTLs (in bold); $2\Delta\ln L$ & $P$ values compare the full model to reduced models in which individual QTLs were eliminated; $n = 139$ .

Position in cM (1.5 LOD interval); LOD score, log odds ratio; Δ*p* LOD*a*, penalized LOD score, change in penalized LOD score compared to the full (best supported) model incorporating all three putative QTLs (in bold); 2*Δ* lnL& *P* values compare the full model to reduced models in which individual QTLs were eliminated; *n* <u>= 139.</u>

### Preference QTL are of large effect

Individually, the measured effect of each of the three QTLs we identified was large, explaining between 23 and 31% of the difference between males of two parental species (Fig. 2B). However, in studies with relatively small sample sizes such as ours (*n* = 139), estimated effects of QTL are routinely over-estimated (a phenomenon known as the “Beavis effect”, after [40]). This is because effect sizes are determined only after significance has been determined, and QTL with artificially high effect sizes – due to variation in sampling – are more likely to achieve ‘significance’.

To determine the extent to which the effects of our QTL may be over-estimated, we simulated QTL across a range of effect sizes, and compared the distribution of measured effects for all simulations to those which would be significant in our analysis (Fig. S3). Our simulations suggest that the reported effects of our QTL are not greatly over-estimated. We first considered what proportion of ‘significant’ simulations would be smaller than our empirically measured effects (Fig. 4A). This suggests that the true effect sizes of our QTL are likely to be large, with somewhat less support for the QTL on chromosome 17. A highly conservative threshold of 95% would suggest that the QTL on chromosome 1 and 18 explain at least 10% and 20% of the difference in behavior between the parental species, respectively. Adopting the median values, our simulations would suggest true effects of 25%, 15% and 30%, or greater, for the QTL on chromosomes 1, 17 and 18, respectively. Given simulated effect sizes similar to those measured empirically, there was little bias among simulation runs that achieved the genome-wide significance threshold (Fig. S3). This suggests that the true effect sizes of our QTL are likely to be large, with somewhat less support for the QTL on chromosome 17.

**Figure 4.**
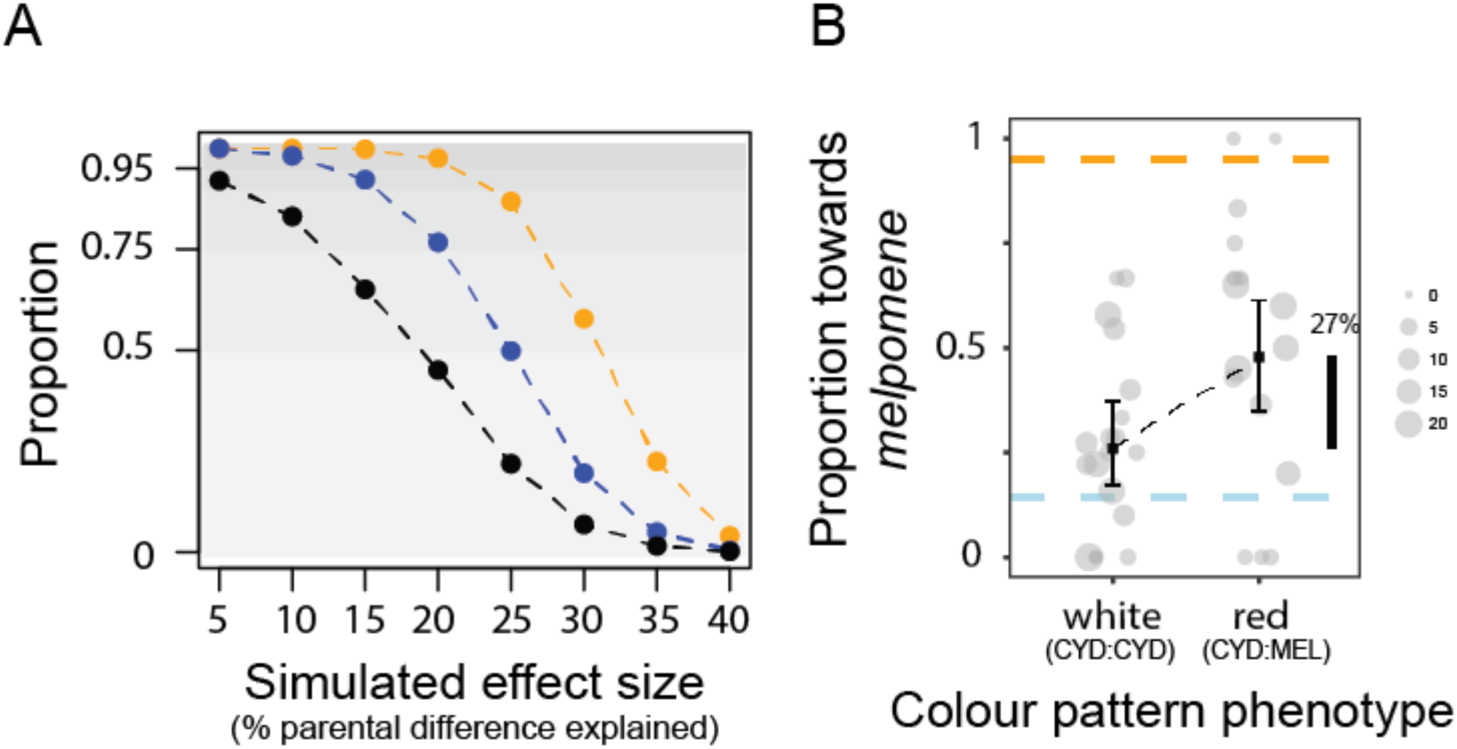
QTL effects in consideration of the to the Beavis effect. A, Proportion of ‘significant’ simulations that would be smaller than our empirically measured effects, for preference QTL on chromosome 1 (blue), chromosome 17 (black), and chromosome 18 (orange). 10000 simulations were run for effect sizes corresponding to between 5 and 40% of the difference in male preference behavior between the parental species. In each case the distribution of sample effect sizes was determined for those simulations that reached the genome-wide significance threshold determined through permutation (Fig. 2). B, Proportion of time males court *H. melpomene* (as opposed to *H. cydno*) females for each of the two genotypes for white (homozygous = CYD:CYD) and red (heterozygous = CYD:MEL) hybrid males for which we were unable to generate RAD data (and so which were not included in our initial QTL analysis). Error bars represent 95% confidence intervals. Lower dashed blue and upper orange bars represent mean phenotypes measured in *H. cydno* and *H. melpomene,* respectively. Circle size depicts total number of ‘courtship minutes’ for each male. Vertical black bars indicate the percentage of the difference measured in the parental species explained.

Although our simulations suggest the effects we have measured are reasonable, ideally we would estimate effect sizes from a population of individuals that were not used to determine significance. In evolutionary biology, follow-up experiments such as this are uncommon; collecting phenotypic data across a large number of hybrid individuals is often a considerable undertaking, and this is similarly true for *Heliconius* behaviors. Nevertheless, we were able to follow-up our results for the QTL on chromosome 18, using a sample of a further 35 backcross males for which preference behavior was measured, but for which we were unable to generate genotype data (and so were not included in our initial QTL analysis). As reported above, the QTL peak (at 0cM) on chromosome 18 is in very tight linkage with the *optix* color pattern locus (at 1.2cM), which controls the presence and absence of the red forewing band. Presence of the red forewing band is dominant over its absence, so that segregation of the red forewing band can be used to perfectly infer genotype at the *optix* locus, even without sequence data. This analysis supports our previous result that the QTL on linkage group 18 is of large effect (Fig. 4B): among these 35 hybrid males, the *optix* locus explains 27% of the difference in behavior between the parental species (*c.f.* 31% for the larger mapping population).

### Admixture is reduced at the preference-color pattern locus on chromosome 18

To consider the effects of major color pattern cue and preference loci on localized gene flow across the genome we used the summary statistic *f*_d_ to quantify admixture between *H. cydno chioneus* and *H. melpomene rosina* (Fig. 3 and S4). *f*_d_ is based on the so-called ABBA-BABA test and provides a normalized measure that approximates the proportional effective migration rate (*i.e. f*_d_ = 0, implies no localized migration of alleles, whereas *f*_d_ = 1, implies complete localized migration of alleles) [41,42]. At the physical location of our behavioral QTL on chromosome 18, which is in tight linkage with the *optix* color pattern locus, there is a substantial reduction in admixture across a ∼1 megabase region. At our other two QTLs, reduced *f*_d_ values (<0.1) are observed for individual 100kb windows associated with all behavioral QTL (specifically, within the 1.5-LOD intervals); but, this is true for many sites across the genome. In addition to mating behavior these two species differ among a number of other behavioral and ecological axes and genomic divergence is highly heterogenous.

### Different preference QTL affect different aspects of behavior

The male preference QTLs we have identified may influence differences in male attraction towards red *H. melpomene* females, or white *H. cydno* females, or towards both female types. To further explore the influence of segregating alleles at these loci we considered the influence of all three QTLs on courtships directed towards each female type separately (Fig. 5). We have already robustly established a significant effect of these loci on variation in the relative amount of time males spent courting each female type (see Fig. 2A). Consequently, although we corrected for multiple testing arising from considering three QTL across the two data sets [37], in these *post-hoc* analyses we did not account for multiple segregating loci across the entire genome (in contrast to the results reported above). This greatly increases our power to detect any influence of the QTLs on attraction towards the two species individually, but also increases the likelihood of false positives. The QTL on chromosome 1 influenced the number of courtships directed towards *H. cydno* females (*F*_1,145_ = 10.85, *P* < 0.01), but had no significant effect on how males behaved towards *H. melpomene* females (*F*_1,145_ = 1.35, *P* > 0.2). In contrast, the QTL on chromosome 17 influenced the degree of courtship directed towards *H. melpomene* (F_1,145_ = 10.08, *P* = 0.011), but not *H. cydno* females (*F*_1,145_ = 0.41, *P* > 0.2). Similarly, the QTL on chromosome 18 had a significant effect on courtships directed towards *H. melpomene* (*F*_1,145_ = 9.93, *P* = 0.012) females (though we note that prior to Bonferroni correction there is also some support for an effect on courtships directed towards *H. cydno* females: *F*_1,145_ = 6.56, *P* = 0.01).

**Figure 5.**
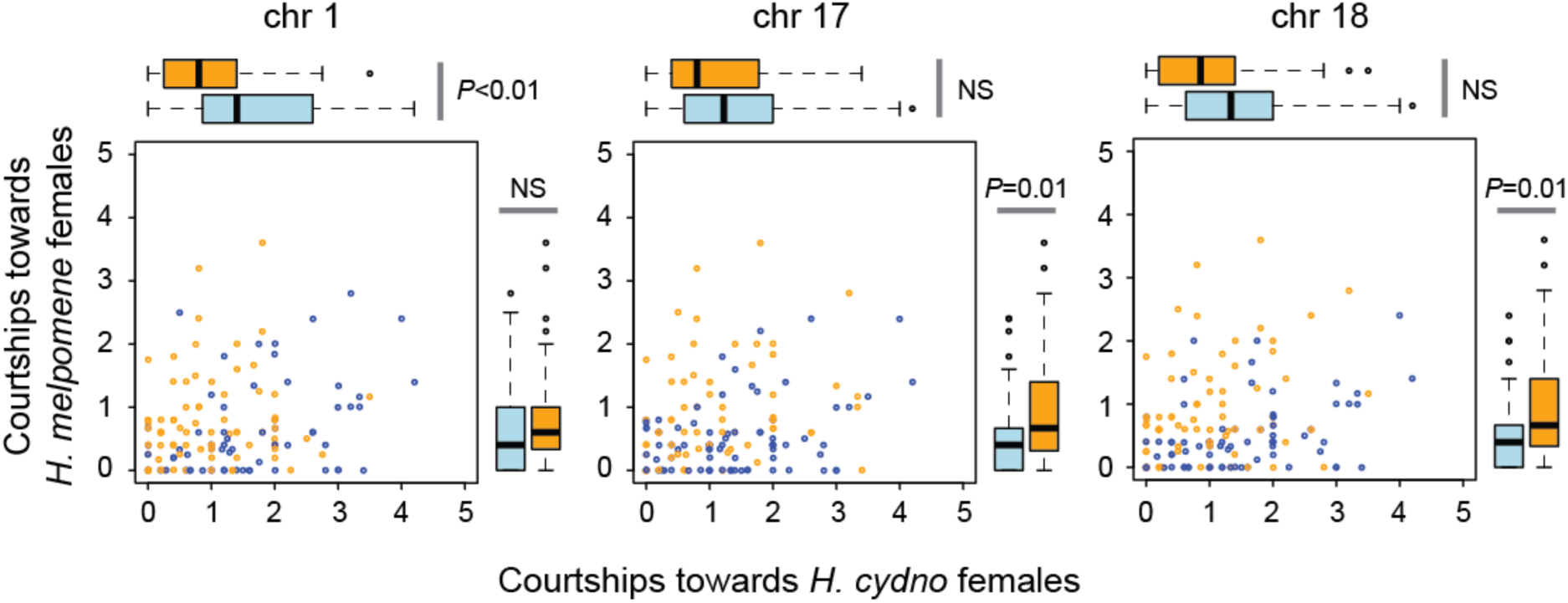
Different QTL affect different aspects of behavior. The QTL on chromosome 1 influences courtship towards *H. cydno,* but not *H. melpomene* females. The opposite is the case for the QTLs on chromosomes 17 and 18, where there is little evidence that either QTL influence courtships directed towards *H. cydno* females. Data presented are for number courtship events corrected by the total number of trials. Blue circles and boxplots represent data for individuals homozygous at each QTL (i.e. CYD:CYD), orange circles and boxplots represent data for individuals heterozygous at each QTL (i.e. CYD:MEL)

## Discussion

Here, we reveal a genetic architecture that will strengthen genetic associations (*i.e.* LD) between key components of reproductive isolation, and so facilitate ecological speciation in the face of gene flow. Specifically, we demonstrate that just three QTLs are largely responsible for an important component of behavioral isolation between two sympatric species of *Heliconius* butterfly. One of these resides only 1.2 (0-4.8) cM from a major color pattern gene. Our results also suggest that all three preference loci are of large phenotypic effect. Because LD between cue and preference loci will arise as a natural consequence of mating preferences [43], these large effect preference loci will further increase LD between ecological and behavioral components of reproductive isolation. Additional smaller effect loci undoubtedly also contribute to variation in male preference, which we would be unlikely to detect without very large sample sizes (a caveat shared with many QTL studies of ecologically relevant behaviors e.g. [15,44,45]). Regardless, our results suggest that during speciation, divergence between populations in both mating cue and the corresponding preference behaviors can have a surprisingly simple genetic architecture.

By ensuring robust genetic associations between components of reproductive isolation, physical linkage between loci for traits influencing pre-and post-mating isolation is expected to facilitate speciation with gene flow [19]. Two of the behavioral QTL we have identified are situated on chromosomes with major color pattern loci (chromosome 1 includes the *K* locus, and chromosome 18 includes the *optix* locus). Both *optix* and the *K* locus have previously been associated with variation in *Heliconius* courtship behaviors [25,35,36]. Nevertheless, we have not previously been able to robustly estimate the position of QTLs along the chromosome. The QTL we identify on chromosome 1 is not tightly linked to the *K* locus. It remains to be seen whether this QTL underlies the association between male preference behavior and the *K* locus phenotype (a shift between white and yellow color pattern elements) previously observed in crosses between *H. cydno* and *H. pachinus* [35], and within a polymorphic population of *H. cydno* [36]. (Although the *K* locus phenotype segregating in crosses between *H. cydno* and *H. melpomene* [39] has not been mapped, it is very likely that it is the same locus as that observed in *H. cydno* and *H. pachinus*). In contrast, our results reveal that the QTL for male attraction on chromosome 18 is tightly linked to the *optix* locus, which controls presence/absence of a red forewing. The mechanistic basis for linkage of trait and preference loci remains unclear. There is no evidence for an inversion at this locus [39]; it also seems unlikely that the same mutations control both wing pattern and the corresponding attraction behavior. However, *optix* is known to function during eye and neural development in *Drosophila* [46], and is expressed in the optic lobe and medulla of pupal *Heliconius* [47], so it is plausible – if unlikely [48] – that the two traits could be controlled by different regulatory elements of the same gene.

Our work joins a small collection of studies in animals where physical linkage is reported to couple loci contributing to preference behaviors and ecological barriers [15,25,35,36,49], as predicted by Felsenstein [19]; and more broadly between loci for cue and preference between incipient species [50–55]. In a seminal study, published almost 20 years ago, Hawthorn and Via [49], showed that QTL for preference and performance for different host plants co-segregate in pea aphids. These insects mate on their host providing a rapid path to speciation. The resolution of molecular markers available at the time allowed linkage to be confirmed to no more than ∼10cM, but even this could substantially impede the break-down of LD: whereas LD between unlinked loci declines by 50% in one generation of random mating, LD between two loci that are 10cM apart would decline by only ∼9% per generation (cyclical parthenogenesis would further reduce recombination in these aphid species). Extending the same logic to our results, LD between the preference locus and *optix* on chromosome 18 would be expected to decline by 1.2 (0-4.6) % per generation (Fig S5), assuming random mating. However, alleles at the behavioral locus result in a preference for the trait controlled by *optix:* LD will be further maintained by non-random mating because warning pattern is a magic trait. As such LD is likely to decline much more slowly than this simple model would suggest.

More recently, Bay and colleagues [15] have reported widespread physical linkage between loci for divergent mate choice and ecological phenotypes in benthic and limnetic populations of three-spine sticklebacks. Two lines evidence support this. First, individual QTLs for mate choice and morphology map to chromosome 14. Second, a polygenic QTL model predicting hybrid position along the benthic-limnetic morphological axis, generated by a previous study [10], explains a significant proportion of variance in mate choice, consistent with physical linkage of ecological and mate choice loci. Our results complement this previous work by explicitly demonstrating tight linkage between assortative mating and ecological traits. In addition, our study shows a much simpler genetic architecture, which should further facilitate the maintenance of LD between traits and which is predicted to facilitate speciation [2].

When mate choice is based on a preference for divergent ecological traits, this will inevitably couple ecological and behavioral components of reproductive isolation. Furthermore, the strength of LD generated will be proportional to the strength of the mating preference, so a genetic architecture with large-effect loci controlling assortative mating will generate stronger LD than a more polygenic architecture. Both our simulations and replication analysis support the existence of large effect QTLs controlling an important interspecific difference in preference behavior. Even if we adopt an especially cautious approach, the QTLs on chromosomes 1 and 18 would explain *at least* 10% and 20% of the difference in male preference behavior, respectively. However, our follow-up analysis, exploiting individuals that were not used to determine significance (thereby evading the Beavis effect), suggests that these estimates are overly conservative; these data explicitly reinforce our initial estimate for the QTL on chromosome 18, which explains ∼30% of the difference between parents. One potential caveat is that the position of the putative QTL and that of *optix* are not the same, but 1.2cM apart; however, any recombination between these loci in the individuals tested will be rare (we expect just 0.42 recombination events between these two loci across 35 individuals), and likely has very limited impact on our estimates of effect size.

We observed a dramatic reduction in admixture (estimated using *f*_d_) at the proximal end of chromosome 18, and specifically on the distal side of *optix* coincident with our QTL. It is tempting to ascribe this to the combined effects of the major preference locus we have identified and the color pattern gene *optix.* However, in the populations studied here, the phenotypic effect of *optix* is more striking than the other color pattern loci, and selection against introgression is likely be stronger at this locus. As a result, tight linkage with *optix* makes it impossible to determine any effects of the preference locus alone. Similarly, it is difficult to infer a signal of reduced admixture due to the behavioral QTLs on chromosomes 1 and 17. Levels of *F*_st_ are high across the genome between *H. cydno* and *H. melpomene* and patterns of admixture across the genome suggest widespread selection against introgression [42]. At this point, the patterns of divergence between *H. cydno* and *H. melpomene* are so heterogenous, it is difficult to disentangle the many processes that could be driving reduced admixture.

A general caveat of our results, alongside other studies of the genetics of assortative mating in *Heliconius* [35,36] and elsewhere (e.g. [15,56]), is that it is hard to distinguish between loci affecting preference behaviors *per se*, from other traits that influence the behavior of the opposite sex. Here, we measured the time *Heliconius* males spend courting a particular female, which may depend not only on male attraction, but also on the female’s response to male behavior (and in turn the male’s response to the female’s behavior). Recent work suggests that *H. cydno* females respond differently to *H. pachinus* than conspecifics males [27]. Although there is currently no evidence that female *Heliconius* use color pattern as an interspecific mating (or rejection) cue (but see [57]), it is not inconceivable, and this could perhaps account for the apparent linkage between male interest and forewing color observed in our study and elsewhere [35,36]. In addition, it is possible that either of these QTLs we identified might influence male pheromones, which has been shown to influence female acceptance behaviors within *H. melpomene* [58]. Nevertheless, using the same hybrids as studied here, we previously demonstrated that individuals that have inherited the red band allele from *H. melpomene* are more likely to court artificial females with the red *melpomene* pattern, implying that the QTL on chromosome 18, at least, influences male response to a visual cue [25]. Regardless of the exact proximate mechanisms involved, the QTLs we identify here influence an important component of male assortative mating behavior.

Overall, the scenario we describe reflects one that modeling broadly predicts will generate a strong overall barrier to gene flow through reinforcement [4]: Specifically, the effects of barrier loci on prezygotic isolation are strong, recombination between pre-and post-mating isolation barrier loci is reduced, and hybridization imposes high costs. Indeed, experimental evidence shows that non-mimetic hybrids between *H. melpomene* and *H. cydno* suffer not only increased predation [24], but also reduced mating success [22] and fertility [59]. In addition, males make a considerable reproductive investment by donating a nutrient-rich spermatophore during mating [60,61], so indirect selection against poorly adapted hybrids could strengthen divergent male preferences. Consistent with a role of reinforcement, *H. melpomene* males from French Guiana, outside the range of *H. cydno*, are less choosy than males from Panama, where the species co-occur and are known to occasionally hybridize [21]; and similar patterns of reproductive character displacement have been observed elsewhere in the *melpomene*-*cydno* clade [62].

Reinforcement is further promoted when indirect selection, resulting from coupling of prezygotic and postzygotic barrier effects, is supplemented by direct selection [4,63]. In *Heliconius,* divergence in male preferences is likely initiated by divergence in wing pattern, and male preferences are observed between populations with few opportunities for hybridization e.g. [64]. Female re-mating is a rare event [65], and males must compete to find virgin females within a visually complex environment [26]. Divergence in female (and male) wing patterns is driven primarily by strong selection for mimicry, and is likely to impose divergent sexual selection on male preferences to improve their ability to find receptive females. This is similar to examples of assortative mating driven by sensory drive, such as in cichlid fishes [66], but it is perhaps less well appreciated that morphological traits under ecological selection (such as *Heliconius* wing patterns) might impose divergent sexual selection on male preferences in a similar fashion.

In addition to a simple genetic architecture, different QTLs appear to control different aspects of preference behavior. Our *post-hoc* analyses suggest that differences associated with QTL1 and QTL17 in the relative amount of time spent courting each female type are driven by differences in attraction to either *H. cydno* or *H. melpomene*, respectively, rather than both species. QTL18 also seems to influence attraction to *H. melpomene* much more strongly than to *H. cydno* females. This genetic modularity, where discrete, independently segregating loci appear to affect different aspects of behavior, may facilitate evolutionary change and innovation by providing a route for rapid evolution of novel behavioral phenotypes [44,67]. In *Heliconius*, this might allow different aspects of mating behavior to evolve independently. It might also allow novel composite behavioral preferences to arise through hybridization and recombination. There is some evidence that this has occurred during hybrid speciation in *Heliconius*. The wing pattern of the hybrid species *H. heurippa* includes both red and yellow pattern elements, which are believed to have originated from the putative parental species *H. melpomene* and *H. cydno,* respectively (local Colombian races of *H. cydno* have a yellow, as opposed to white, forewing band) [23]. Not only do *H. heurippa* males prefer this combined pattern over the ‘pure’ red or yellow patterns of *H. melpomene* and *H. cydno* [23], but ‘recreated *H. heurippa*’, obtained in first generation backcrosses between *H. melpomene* and *H. cydno*, prefer the pattern of *H. heurippa* over that of the two putative parents [68]. This is consistent with a hypothesis in which introgression and subsequent recombination of preference alleles are responsible for novel behavioral phenotypes, although further work would be needed to confirm this.

In conclusion, the genetic architecture we demonstrate here will promote the evolution of behavioral isolation by strengthening genetic associations between cue and preference. Disassociation of alleles at loci that are physically close on the chromosome is slower compared to that between alleles at more distant loci (due to reduced crossing over), or at loci on different chromosomes. Similarly, the substitution of large effect alleles will also increase linkage disequilibrium between cue and preference, even if they are not physically linked, because preference alleles of larger effect will more often find themselves in the same genome as alleles for the corresponding cue, compared to preference alleles with smaller effects. We cannot currently distinguish whether preference QTL result from single adaptive mutations, or represent multiple functional loci that have built up during the course of speciation. Nevertheless, the genetic basis of *Heliconius* mate preferences is remarkably similar to that for differences in the wing pattern cue. Differences in individual color pattern elements probably do involve multiple, sequential mutations (which target the same gene(s)), but ‘ready-made’ alleles of large phenotypic effect can be brought together in new combinations through adaptive introgression. The existence of large effect preference loci, potentially influencing different aspects of behavior, could similarly facilitate the origin of novel phenotypes through introgression, and further facilitate rapid speciation.

## Methods

### Butterfly collection, rearing and crossing design

All butterfly rearing, genetic crosses and behavioral experiments were conducted at the Smithsonian Tropical Research Institute in Panama between August 2007 and August 2009. We collected wild *Heliconius cydno chioneus* and *Heliconius melpomene rosina* from Gamboa (9°7.4’N, 79°42.2’ W, elevation 60 m) and the nearby Soberania National Park, Panama. These were used to establish stocks maintained in insectaries in Gamboa, which were further supplemented with wild individuals throughout the experimental period. We established interspecific crosses by mating wild caught *H. melpomene* males to *H. cydno* females from our stock population. In interspecific crosses between *Heliconius cydno* females and *Heliconius melpomene* males, F1 hybrid females are sterile, restricting us to a backcrossing design. We generated backcross broods to *H. cydno* and *H. melpomene* by mating F1 males to virgin females from our stock populations. Brood mothers were kept individually in cages (approx. 1 or 2 x 2x2m), and provided with ∼10% sugar solution, a source of pollen and *Passiflora* shoots for oviposition. Eggs were collected daily and caterpillars raised individually in small pots until 4th or 5th instar, and then either in groups or individually until pupation. Caterpillars were provided with fresh *Passiflora* leaves and shoots daily.

### Behavioral assays

We measured male attraction to *H. melpomene* and *H. cydno* females in standardized choice trials [25,38]. Males were allowed to mature for at least 5 days after eclosion before testing. Males were introduced into outdoor experimental cages (1x1x2m) with a virgin female of each species (0 – 10 days matched for age). Fifteen-minute trials were divided into 1 min intervals, which were scored for courtship (sustained hovering or chasing) directed towards each female as having occurred or not occurred. Accordingly, if a male courted the same female twice within a minute interval, it was recorded only once; if courtship continued into a second minute, it was recorded twice. Where possible, trials were repeated for each male (median = 5 trials). From these trials we generated a large dataset used in subsequent analyses which includes the total number of ‘courtship minutes’ directed towards *H. melpomene* and the number of ‘courtship minutes’ *H. cydno* females (Table S2). The QTL analysis considered the proportion of total ‘courtship minutes’ directed towards *H. melpomene, i.e.* ‘courtship minutes’ directed towards *H. melpomene /* (‘courtship minutes’ directed towards *H. melpomene +* ‘courtship minutes’ directed towards *H. cydno*) = “the relative amount of time males spent courting red *H. melpomene* and white *H. cydno* females” = “relative courtship time”. In total we conducted 1347 behavioral trials, and collected data from 28 *H. cydno*, 16 *H. melpomene*, 23 F1 hybrid, 29 backcross-to-*melpomene* hybrid and 196 backcross-to-*cydno* hybrid males (of which 11 performed no courtship behaviors).

### Genotyping and linkage map construction

Genotyping and construction of linkage maps has been described elsewhere [39]. In brief, backcross hybrids and associated parents were preserved in 20% DMSO and 0.25M EDTA (pH 8.0) and stored at-20°C. DNA was extracted with Qiagen DNeasy Blood& Tissue Kits following the manufacture’s protocol for animal tissue. Individuals were genotyped using a RAD-sequencing approach [69] and sequenced by BGI using the Illumina HiSeq 2500. Sequences were then aligned to version 2 of the *H. melpomene* genome [70] using Stampy v1.0.23 [71]. Duplicates were removed with Piccard tools v1.135 (http://broadinstitute.github.io/picard/), and genotype posteriors called using SAMtools v1.2. Interspecific linkage maps were constructed using Lep-MAP2 [72] and modules from Lep-MAP3 as described in [39]. To obtain the genotypic data for QTL mapping, the parental-phased data was obtained using Lep-MAP3 option outputPhasedData=1. This option imputes data based on input genotypes and the map order. These data were then compared to the subset of markers in which grandparents could be used to phase the data for each family and chromosome using custom scripts. Family and chromosome was inverted when required to obtain matching phases. Finally, the non-informative markers between inferred recombinations were masked (i.e. set to missing) to account for the fact the exact recombination position was not known for these regions.

### Data analysis

All QTL analyses were performed on backcross-to-*cydno* hybrid males in which the preference behaviors segregate. We were able to generate genotype data for 146 of the 196 backcross-to-*cydno* hybrid males for which we recorded behaviors in our choice trials. The remaining 50 individuals include males from which we were unable to extract sufficient DNA, were poorly sequenced, or were lost in the insectaries most often due to ants or other predators. For each backcross individual, we calculated the probabilities of the two alternative genotypes at every marker and centiMorgan (cM) position along the chromosomes, conditional on the available marker data, using R/qtl package [73]. R/qtl uses a hidden Markov model to calculate the probabilities of the true underlying genotypes given the observed multipoint marker data. We then tested for an association between phenotype and genotype at each position using generalized linear mixed models (GLMMs) with binomial error structure and logit link function (implemented with the R package lme4). We first considered the relative time males courted *H. melpomene* as opposed to *H. cydno* females. For each position along the genome we modeled the response vector of the number of ‘courtship minutes’ towards *H. melpomene* vs ‘courtship minutes’ towards *H. cydno* with the genotype probability as the independent variable. LOD scores were obtained by comparing this to a null model in which genotype probability was not included. An individual level random factor was included in all models to account for over-dispersion. This approach is analogous to the to the Haley-Knott regression implemented in R/qtl [54, 55], but more appropriately accounts for the non-normal structure of our data and for differences in total courtship data recorded for each individual [56]. Seven individuals were excluded from these analyses for which, although tested in multiple trials, no courtship towards either female type was recorded. Using permutation [57], we determined the genome-wide significance threshold for the association between marker genotype and phenotype (alpha = 0.05, n = 1000 permutations) as LOD = 2.99. By using our GLMM approach we had more power to detect QTL than would be permitted by adopting non-parametric methods. Nevertheless, we repeated all QTL analyses using non-parametric interval mapping in R/qtl, using the ‘scanone’ and ‘model = “np”’ commands. Results of non-parametric analyses are reported in the supplementary materials (Table S2).

To consider all three QTL identified in our initial genome scans together, we again modeled the number of ‘courtship minutes’ towards *H. melpomene* vs ‘courtship minutes’ towards *H. cydno* but with the genotype at the max LOD score for each QTL as explanatory variables. The fully saturated GLMM, including all three pairwise interactions, was simplified in a stepwise manner using likelihood ratio tests, but always retaining individual id as a random factor. To further test for effects of each QTL we compared the penalized LOD scores of the full model (including all three QTL as additive effects) to reduced models in which each QTL was eliminated in turn. The penalized LOD score is calculated as: pLODa(γ) = LOD(γ) –T| γ |, where γ denotes a model, | γ | is the number of QTL in the model and T is a penalty determined through permutation (i.e. the genome-wide significance threshold = 2.99) [58].

Finally, to determine the contribution of each QTL to variation in courtship time towards *H. cydno* and *H. melpomene* females separately, we considered the total number of ‘courtship minutes’ directed to each female type, correcting for the number of trials. We included all 146 backcross males for which we had genotype data in this analysis. We square-root transformed courtship minutes/trial and then used the makeqtl() and fitqtl() functions in R/qtl [54] to determine significance. Model residuals were inspected visually for an approximate normal distribution, and we tested for homogeneity of variance with Levene’s tests (*H. cydno* females: F_3,142_ = 0.27, P > 0.02; *H. melpomene* females: F_3,142_ = 0.39, P > 0.02). We corrected *P* values to account for the 6 tests (i.e. 3 loci x 2 species) [37].

### Simulations

We used simulations to estimate potential inflation of measured effect sizes due to the Beavis effect. We generated 10000 simulated data-sets for each of a range of ‘true’ effect sizes for each significant QTL (i.e. on chromosomes 1, 17 and 18), using the R package simr [74]. For each of these we determined the LOD score and compared it to our genome-wide significance threshold (i.e. LOD = 2.99). This allowed us to compare i) the entire range of simulated effects (where the mean is expected to equal the ‘true’ effect size), with those that would be significant given our sample size (n = 139) and linkage map (figure S1), and ii) the empirically measured effects with simulated effects that would be significant (Fig. 3A).

### Admixture analysis

We investigated heterogeneity in admixture across the genome between *H. melpomene rosina* and *H. cydno chioneus* using *f*_d_, which provides an approximately unbiased estimate of the admixture proportion [41,42]. This analysis made use of available whole genome sequence data for *H. melpomene rosina* (N=10) and *H. cydno chioneus* (N=10) from Panama, with *H. melpomene melpomene* from French Guiana (N=10) serving as the allopatric ‘control’ population and two *Heliconius numata* individuals as outgroups [42].

### Code and data availability

Scripts and raw data used in analyses are available online: DRYAD REPOSITORY XXX. *f*_d_ was computed in 100 kb windows using the python script ABBABABAwindows.py, available from github.com/simonhmartin/genomics_general. Sequence data used to make the linkage maps have previously been submitted to the European Nucleotide Archive (http://www.ebi.ac.uk/ena) [39], accession number ERP018627.

## Acknowledgements

We are grateful to Moisés Abanto, Josephine Dessmann, Janet Scott and Bas van Schooten for help in the insectaries, to Justin Touchon for advice regarding statistics, and to Stephen Montgomery, Markus Möst, Ricardo Pereira, Alexander Hausmann and Vera Warmuth for comments on the manuscript. We are also grateful for insightful referee comments which greatly improved this manuscript. We thank the Smithsonian Tropical Research Institute for support and the Ministerio del Ambiente for permission to collect butterflies in Panama. We are also grateful to Edinburgh Genomics for sequencing support. RMM is funded by a DFG Emmy Noether Fellowship, and was also funded by a Junior Research Fellowship at King’s College, Cambridge and by the ERC grant 339873 Speciation Genetics awarded to CJ. JD was funded by a Herchel Smith Postdoctoral Fellowship. SHM was funded by a research fellowship from St John’s College, Cambridge.

## Supporting Information

**Figure S1.**
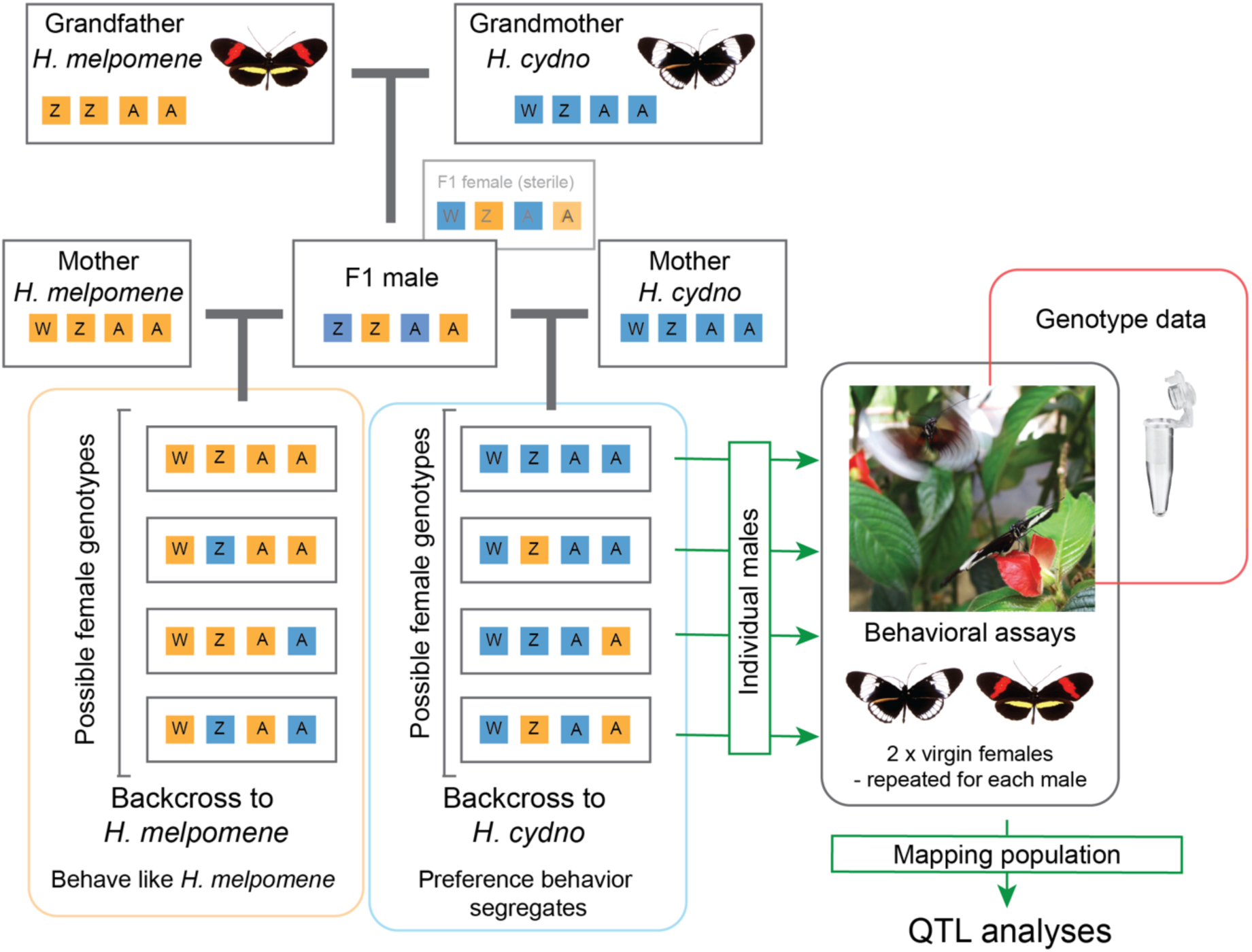
Overview of crossing design. Colored boxes represent segregating *H. cydno* (blue) and *H. melpomene* (orange) alleles; Z and W refer to the alleles on the sex-chromosomes and A to those on autosomes.

**Figure S2.**
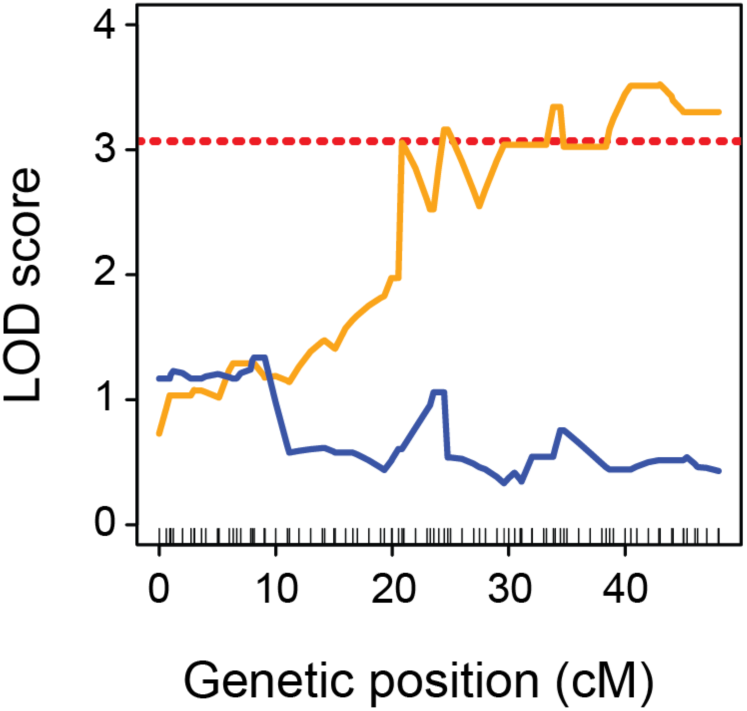
QTL analysis of variation of mate preference for individuals with alternative genotypes at LG18@0cM. QTL associated with the proportion of time males court *H. melpomene* (as opposed to *H. cydno*) females on chromosomes 17 for individuals homozygous (i.e. white, CYD:CYD = blue line) and heterozygous (i.e. red, CYD:MEL = orange line) at LG18@0cM. Dashed line represents log odds ratio (LOD) significance threshold (*i.e.* genome-wide alpha = 0.05) for heterozygous (*i.e.* red, CYD:MEL) individuals. Dashes along the x-axis indicate position of genetic markers (SNPs).

**Figure S3.**
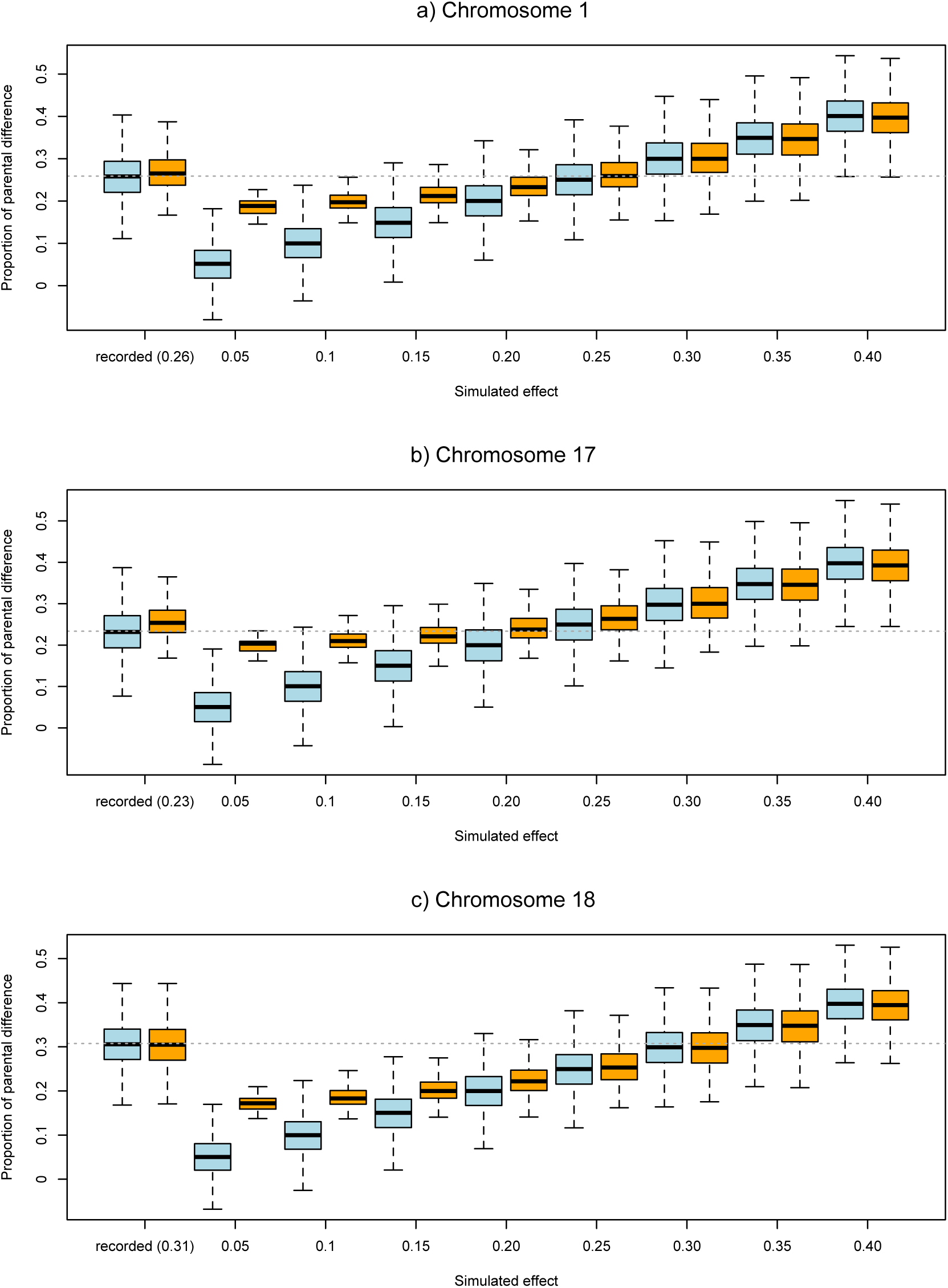
Simulations suggest QTL effect sizes are not greatly overestimated. For each simulated effect size, the distribution of all simulated effects (blue) and those which would be significant in our analysis (*i.e.* LOD ≥ 2.99) (orange) are shown. In each case, ‘recorded’ refers to the empirically measure effect size.

**Figure S5.**
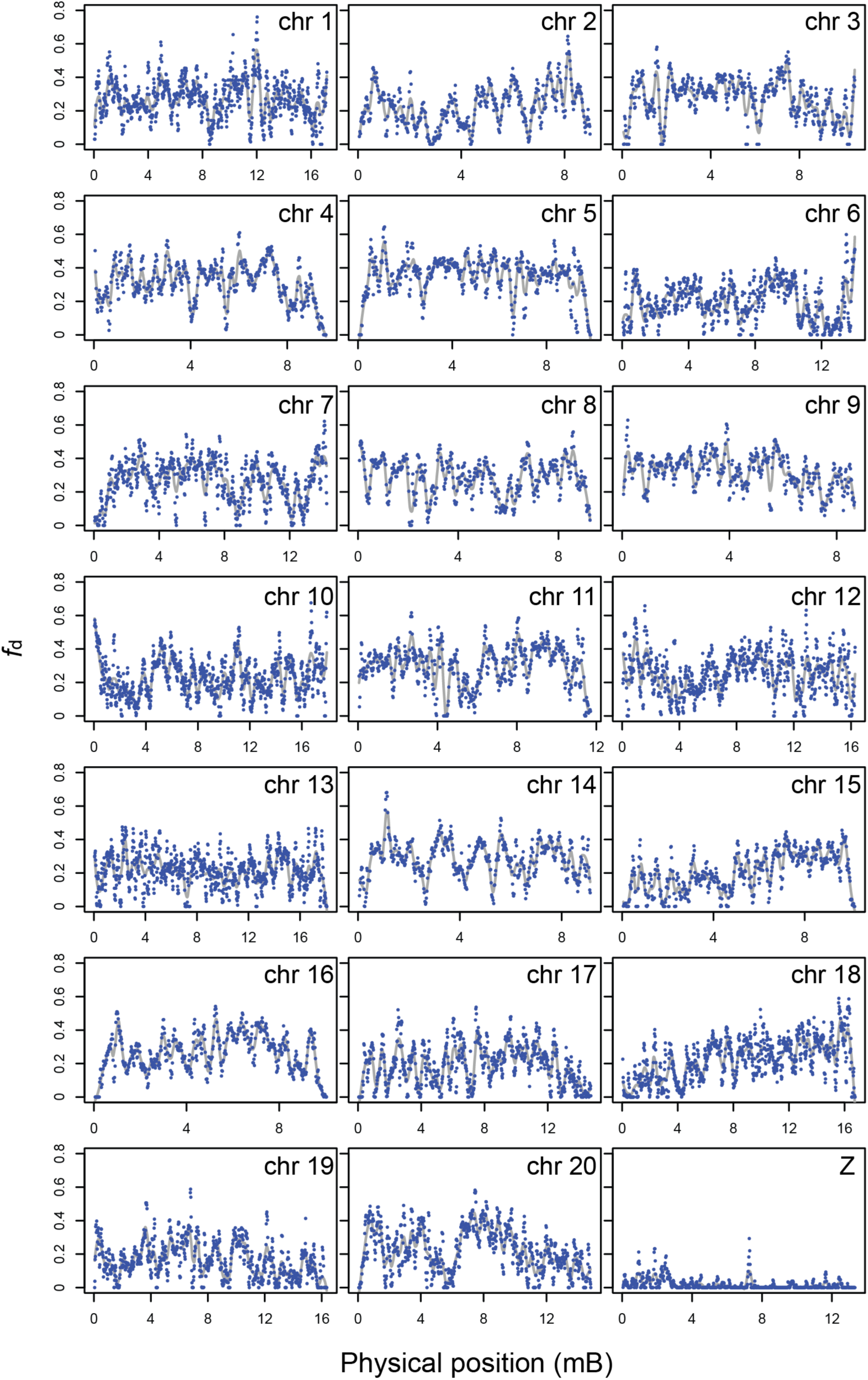
Localized levels of admixture (*f*_d_) across all 21 chromosomes. Blue points represent *f*_d_ values for 100kb windows. *f*_d_ was measured between *H. melpomene rosina* and *H. cydno chioneus* individuals.

**Figure S6.**
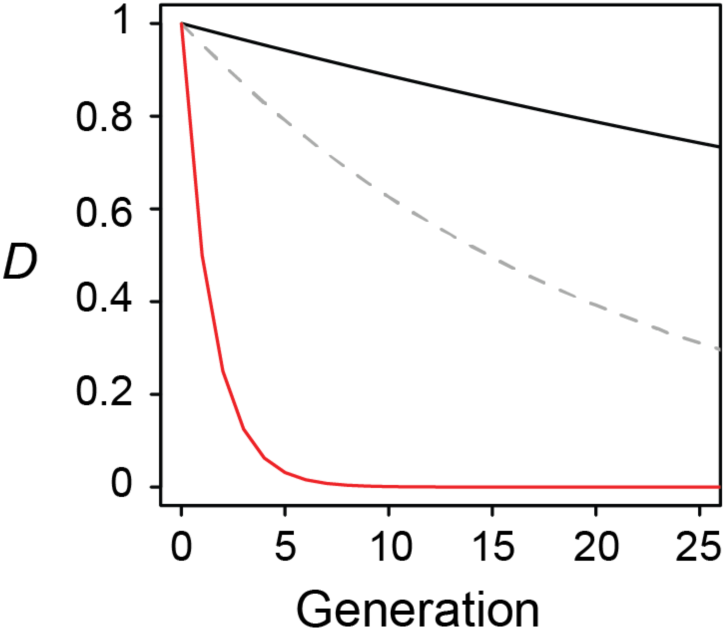
Decline in linkage disequilibrium (LD) of linked and unlinked loci under an assumption of random mating. Whereas linkage disequilibrium (*D*) between unlinked loci (red solid line) declines by 50% in one generation of random mating, LD between two loci that are 1.2cM apart would decline by only 1.2 % per generation (black solid line), and LD between two loci that are 4.8cM apart (gray dashed line) would decline by only 4.6% per generation.

**Table S1.** Summary of genome-wide QTL analyses using binomial GLMM methods (reported in main text) and non-parametric methods implemented in R/qtl. Position in cM refers to the position the peak lod score, i.e. the most likely genetic position of the putative QTL, for each QTL. *P* values are determined through permutation as described in the methods.

| Binomial GLMM |  |  |  | Non-parametric |  |  |
| --- | --- | --- | --- | --- | --- | --- |
| Chromosome | Position (cM) | LOD | $P$ | Position (cM) | LOD | $P$ |
| 1 | 4.23 | 4.54 | <0.001 | 4.23 | 3.6 | ~0.009 |
| 17 | 24.47 | 3.5 | ~0.013 | 24.47 | 2.89 | ~0.049 |
| 18 | 0 | 6.83 | <0.001 | 0 | 5.34 | <0.001 |

**Table S2.Courtship data and total trials for all 292 individuals included in the study.**Type CYD = pure *H. cydno chioneus*; MEL = pure *H. melpomene rosina*; F1 = first generation hybrids (*H. cydno chioneus* mother and *H. melpomene rosina* father); BC = backcross to *H. cydno chioneus*; and BM = backcross to *H. melpomene rosina.*

*See attached.csv file:* Table_S2.csv

